# A highly active *Burkholderia* polyketoacyl-CoA thiolase for production of triacetic acid lactone

**DOI:** 10.1101/2022.12.04.519061

**Authors:** Zilong Wang, Seokjung Cheong, Jose Henrique Pereira, Jinho Kim, Andy DeGiovanni, Yifan Guo, Guangxu Lan, Carolina Araujo Barcelos, Robert Haushalter, Taek Soon Lee, Paul D. Adams, Jay D. Keasling

**Affiliations:** Joint BioEnergy Institute, Emeryville, California 94608, USA; Department of Chemical and Biomolecular Engineering, University of California, Berkeley, California 94720, USA; Biological Systems and Engineering Division, Lawrence Berkeley National Laboratory, Berkeley, California 94720 USA; Molecular Biophysics & Integrated Bioimaging Division, Lawrence Berkeley National Laboratory, Berkeley, CA, USA; Department of Bioengineering, University of California, Berkeley, Berkeley, CA 94720, USA; Advanced Biofuels & Bioproducts Process Demonstration Unit, Lawrence Berkeley National Laboratory, Berkeley, CA, 94720, United States; QB3 Institute, University of California, Berkeley, California 94720, USA; Center for Biosustainability, Danish Technical University, Lyngby, DK; Center for Synthetic Biochemistry, Institute for Synthetic Biology, Shenzhen Institutes of Advanced Technologies, Shenzhen 518055, China

**Author notes:** These authors contributed equally to this study (Zilong Wang, Seokjung Cheong).

## Abstract

Triacetic acid lactone (TAL) is a platform chemical biosynthesized primarily through decarboxylative Claisen condensation by type III polyketide synthase 2-pyrone synthase (2-PS). However, this reaction suffers from intrinsic energy inefficiency and feedback inhibition by and competition for malonyl-CoA. TAL production through non-decarboxylative Claisen condensation by polyketoacyl-CoA thiolase alleviates many of these disadvantages. We discovered five more thiolases with TAL production activity by exploring homologs of a previously reported polyketoacyl-CoA thiolase, BktB, from *Cupriavidus necator*. Among them, the BktB homolog from *Burkholderia* sp. RF2-non_BP3 has ∼ 30 times higher *in vitro* and *in vivo* TAL production activity and led to ∼10 times higher TAL titer than 2-PS when expressed in *Escherichia coli*, achieving a titer of 2.8 g/L in fed-batch fermentations. This discovery of a novel polyketoacyl-CoA thiolase with superior TAL production activity paves the way for realization of total biomanufacturing of TAL.

## Introduction

Microbial biosynthesis of chemicals and fuels is an important research topic as it seeks to replace petroleum feedstocks with renewable biomass or one-carbon gasses, reduce the emissions of greenhouse gasses and pollutants, and improve flexibility and capital efficiency ^1–3^. Enzymatic reactions constitute the biosynthetic pathways to those molecules, so the selection of enzymes with high activity and selectivity towards the target product is crucial to improve the titer, rate, and yield (TRY). The rise of bioinformatics and systems biology has offered more diverse tools for enzyme discovery and selection^2,4,5^, and directed evolution and computational- and artificial intelligence-assisted rational design can be used to further enhance the desired enzyme activity and performance^3,6–9^.

Triacetic acid lactone (4-hydroxy-6-methyl-2-pyrone, TAL) is a platform chemical used in the synthesis of food preservatives and additives, fragrances, and a variety of other chemicals^10,11^ and polymers^12–14^. As such, it is a privileged molecule, and it can be synthesized both chemically and biologically^15^. Plants such as *Gerbera hybrida* (Daisy) naturally produce TAL through two rounds of decarboxylative Claisen condensation with acetyl-CoA as initial precursor and malonyl-CoA as extension units catalyzed by a type III polyketide synthase, 2-pyrone synthase (2-PS) (**Fig. 1a**)^16^. Heterologous expression of 2-PS enabled TAL synthesis from different carbon sources in several platform microbial hosts like *Escherichia coli*^17–19^, *Saccharomyces cerevisiae*^17,20–22^, *Yarrowia lipolytica*^14,23,24^ and *Rhodotorula toruloides*^25^, with some publications reporting titers of more than 20 g/L. However, use of 2-PS and malonyl-CoA has several disadvantages: ATP consumption to produce malonyl-CoA from acetyl-CoA by acetyl-CoA carboxylase (ACC); allosteric inhibition of malonyl-CoA on ACC; and competition for malonyl-CoA with native pathways like fatty acid biosynthesis^26^. In 2020, Tan et al. discovered that BktB from *Cupriavidus necator* is a polyketoacyl-CoA thiolase and demonstrated its activity to produce polyketides, including TAL, through repetitive non-decarboxylative Claisen condensation reactions with acetyl-CoA as both precursor and extension unit *in vitro* or *in vivo* in *E. coli* (**Fig. 1a**)^27^. The omission of malonyl-CoA by polyketoacyl-CoA thiolase circumvents the disadvantages of 2-PS, so polyketoacyl-CoA thiolase is a promising alternative enzyme for microbial biosynthesis of TAL.

**Fig. 1.**
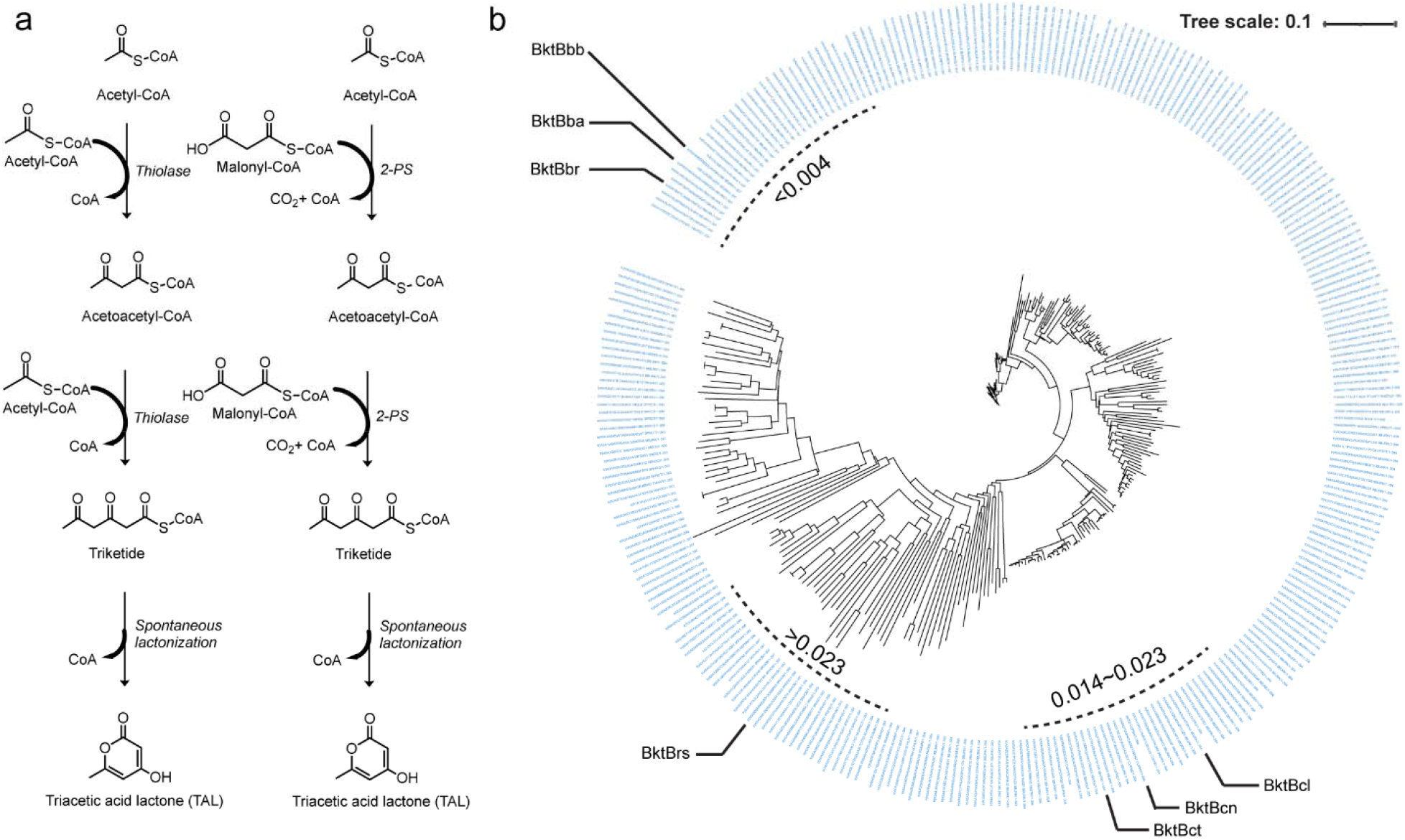
Biosynthetic pathways for triacetic acid lactone and evolutionary analysis of BktBcn homologs. a) Two pathways for TAL production catalyzed by BktB and 2-PS. b) Circular phylogenetic tree constructed after MAFFT alignment with 1798 homologous sequences, of which α-helix-3 and α-helix-5 were identified for the alignment. Other regions of proteins were excluded from the alignment, resulting in a specific focus on the featured region of homologs functionally equivalent to BktBcn.

Here, we report a BktB homolog from *Burkholderia* sp. RF2-non_BP3 (bktBbr, WP_059700748.1), discovered through tests on candidates selected from thousands of BktB homolog pools with different phylogenetic distances to BktB from *Cupriavidus necator* (BktBcn). In addition to possessing the advantages of being a polyketoacyl-CoA thiolase compared to 2-PS, it also has ∼30 times higher TAL-producing activity *in vitro* and *in vivo* than BktBcn and ∼10 times higher activity than 2-PS. Expression of *bktBbr* in *E. coli* led to TAL titers in excess of 2.8 g/L in a fed-batch bioreactor, the highest reported TAL titer from *E. coli*. We anticipate that its implementation in other microbial hosts more tolerant than *E. coli* to TAL, such as yeasts, could further increase the titer.

## Results

### Discovery of novel BktB homologs for triacetic acid lactone production

Unlike most other thiolases, BktBcn is a polyketoacyl-CoA thiolase able to accept acetoacetyl-CoA as the substrate for its non-decarboxylative Claisen condensation, generating TAL through spontaneous cyclization of the triketide CoA. Hereafter, for convenience, all thiolase homologs of BktBcn will be termed “BktB”s. In the first step, the protein sequence of BktBcn was aligned using MAFTT with 2910 thiolase sequences obtained from the Uniprot database. Based on the previous findings about *C. necator* BktB structure (PDB, 4W61) possessing unique features in α-helix-3 and α-helix-5 that expand the substrate binding pocket relative to PhbAzr ^28^, an additional analysis focusing on the α-helix-3 and α-helix-5 was performed, which filtered the sequence pool from 2910 to 1798. Finally, a circular version of a phylogenetic tree with the top 300 sequences was constructed accordingly by EMBL-EBI Phylogeny (**Fig. 1b**). Hereafter, the name of BktBs from various microbial species are simplified as BktBxx, with xx representing the genus and species names, respectively (**Extended Data Table 1**). Six BktB candidates were then selected based on the evolutionary distance in three clusters, <0.004, 0.014∼0.023 and >0.023. Admittedly, there are still many homologs that are not addressed in this study, but it does not influence our aim of selecting different homologs at distinct evolutionary distances.

To validate if the selected BktB candidates could produce TAL using acetoacetyl-CoA, all seven BktB proteins including BktBcn were expressed in *E. coli* BL21(DE3) and purified for *in vitro* studies (**Extended Data Fig. 1a, 1b**). Except for the nearly insoluble BktBrs, which failed to purify (**Extended Data Fig. 1a, 1b**), all other BktB candidates converted acetyl-CoA and acetoacetyl-CoA into TAL (**Fig. 2b, Extended Data Fig. 1c**), leading to a hypothesis that BktB TAL-producing activity exists quite broadly in thiolases. LC-MS analysis revealed that BktBs from *Burkholderia* species produced TAL at a faster rate than BktBs from other microorganisms.

**Fig. 2.**
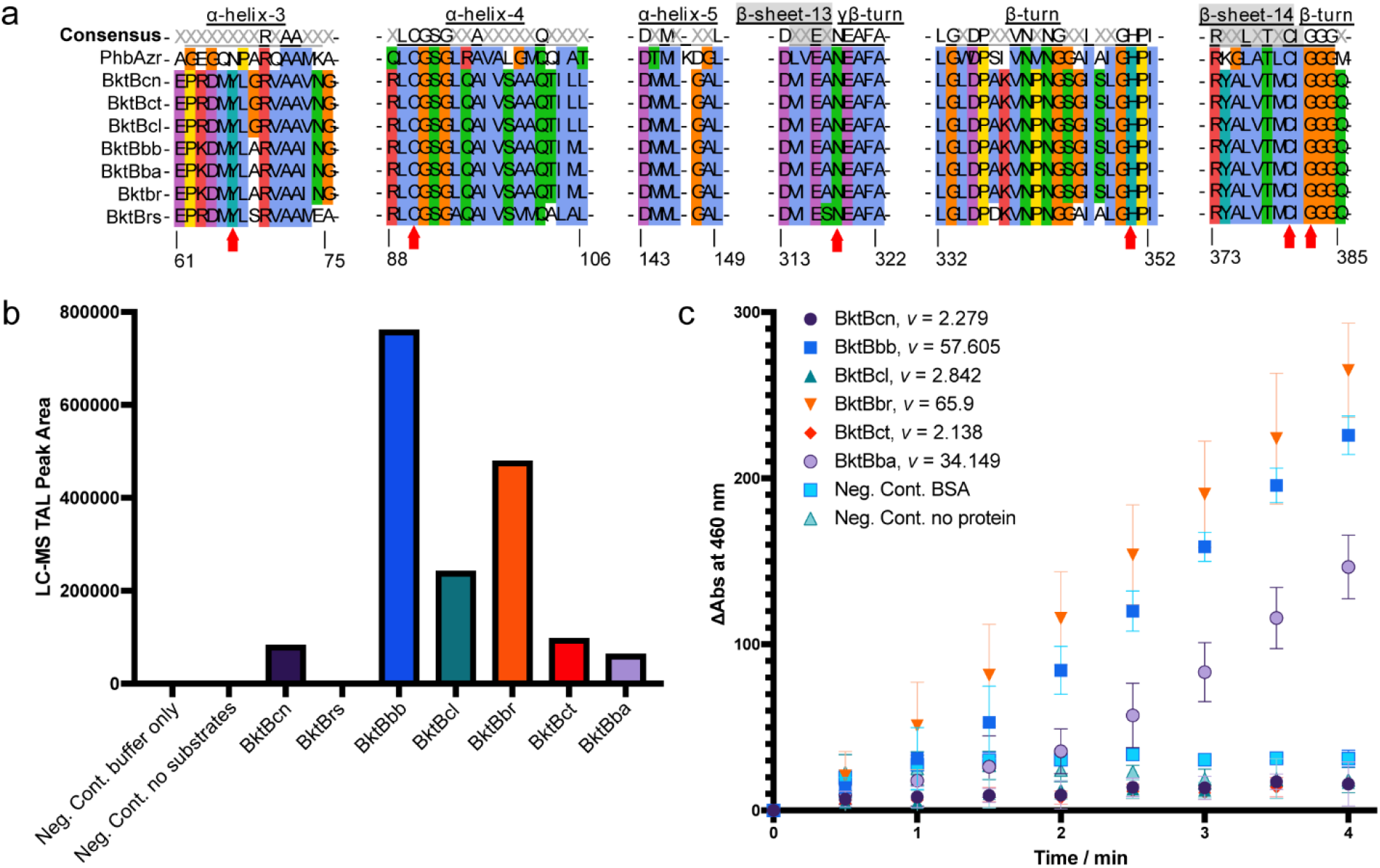
Protein sequence alignment of selected BktB proteins and the kinetics under different conditions. a) Regions of importance including the featured Y66 and α-helix-5, active sites of the sequences are highlighted. PhbAzr is used to show the difference between classic Thiolase II and BktBs. Red arrows indicate amino acids in the active site. X represents the non-consensus amino acids in the alignment. The α-helices, β-sheets and β-turns are underlined, and the β-sheet has a gray background in the consensus row. Amino acid positions referred to positions in BktBcn. b) Functional determination of TAL formation by LC-MS with various BktBs *in vitro*.The reaction contains 1 mM acetyl-CoA and 1 mM acetoacetyl-CoA, incubated at room temperature for 30 min. c) Enzyme assays of six soluble BktBs. The reaction contains 0.1 mM acetyl-CoA and 0.1 mM acetoacetyl-CoA, monitored at 460 nm absorbance of NADH formation coupled with α-KGDH reaction at room temperature. Rate of enzyme reactions is indicated in the legend. BSA is used as the non-relevant protein for the negative control.

### *Burkholderia* BktBs displaying superior enzyme activity

To better understand how the seven selected BktBs are different from each other, protein sequences and enzyme kinetics were analyzed. With the known structures of thiolase PhbAzr from *Z. ramigera* (PDBs 1DLV and 1DM3) and BktBcn (PDB 4W61), accurate secondary structures of the entire amino acid sequence can be depicted. As reported, Y66 is a unique feature of BktBs^28^ compared with thiolase PhbAzr, which shares the ability to condense two units of acetyl-CoA into acetoacetyl-CoA with BktBcn^29^ but cannot synthesize TAL^30^. Important regions in the alignment of seven BktB protein sequences and PhbAzr highlight that these proteins have identical active sites of C90, N317, H350, C380 and G382 (amino acid positions referred to positions in BktBcn), with the conserved α-helix-5 for thiolase biosynthetic activity. The difference in α-helix-3 diverges PhbAzr from BktBs, of which PhbAzr has a shorter helix with the amino acid sequence PARQAAMK, possibly as well as the variation in the loop region of β-turn around the active site residue H350. Notably, *Burkholderia* BktBs are almost identical in the highlighted regions with BktBcn, except for A68, which is only consistent in *Burkholderia* BktBs but not in all other BktBs (**Fig. 2a**). Combining this with the aforementioned LC-MS *in vitro* TAL biosynthesis data showing that *Burkholderia* BktBs generate more TAL than other BktBs (**Fig. 2b**), we hypothesize that *Burkholderia* BktBs are more active in enzyme activity.

There are two common ways to determine the enzyme activity, either monitoring the decrease of the substrates or the increase of the products. Monitoring at UV wavelength 298 nm has been widely used for measuring the kinetics of TAL producing enzymes, including 2-PS and BktB^18,27^. Here, we determined the kinetics of TAL formation by the consumption of CoA substrates, coupled with NADH formation by α-ketoglutarate dehydrogenase (α-KGDH) utilizing free CoA released during TAL formation. Monitoring NADH formation by excitation at 360 nm and emission at 460 nm is more sensitive than directly monitoring TAL absorbance at 298 nm, which overlaps with protein absorbance at 280 nm. The activities of all six soluble BktBs were compared by reacting with 0.1 mM acetyl-CoA and 0.1 mM acetoacetyl-CoA. All three *Burkholderia* BktBs exhibited much higher activity than all other BktBs, especially the fastest *Burk* sp. RF2-nonBP3 BktB activity, which was surprisingly nearly 30-fold higher than BktBcn (**Fig. 2c**). As such, *Burkholderia* BktBs should be very efficient in TAL production *in vivo*.

Genes encoding the tested BktBs along with 2-PS and the negative RFP control were then cloned into pBbA5a^31^, a medium-copy BioBrick vector with medium strength promoter, and the resulting vectors were transformed into *E. coli* JBEI-3695 (BW25113 Δ*adhE* Δ*ldhA* Δ*frdBC* Δ*pta*). The *E. coli* strains harboring these resultant plasmids were grown, and the TAL concentration was measured. JBEI-3695 was used as the host strain, as its deficiency in most of mixed-acid fermentation enzymes improved the supply of substrate acetyl-CoA, which is converted to another substrate acetoacetyl-CoA by native acetyl-CoA acetyltransferase AtoB or overexpressed BktBs themselves (**Fig. 3a**). After 48-hour post-induction growth in 24-well plates under different temperatures, we determined that 25 oC growth led to highest TAL titers **(Extended Data Fig. 3)**. Strains expressing *Burkholderia* BktBs produced more than 75 mg/L TAL, while most other BktBs only produced approximately 10 mg/L titer. The titer of the strain overexpressing 2-PS produced approximately 30 mg/L, and strains with RFP and BktBrs had no detectable TAL production. Among three *Burkholderia* BktBs, BktBbr yielded 0.3 g/L TAL, 30 times the titer from BktBcn, resembling their *in vitro* test results (**Fig. 3b**). BioBrick vectors with different copy numbers and promoter strengths were then tested for BktBbr, and as expected, the TAL titer improved when the vector copy number was higher. Further implementation of a stronger promoter on pBbE1a resulted in 0.8 g/L TAL production (**Fig. 3c**). JBEI-3695 pBbE1a-bktBbr, the strain with highest TAL production in 24-well plates, was also grown in shake flasks with glycerol and glucose as carbon sources, resulting in 0.4 g/L and 0.2 g/L of TAL, respectively **(Fig. 3d)**.

**Fig. 3.**
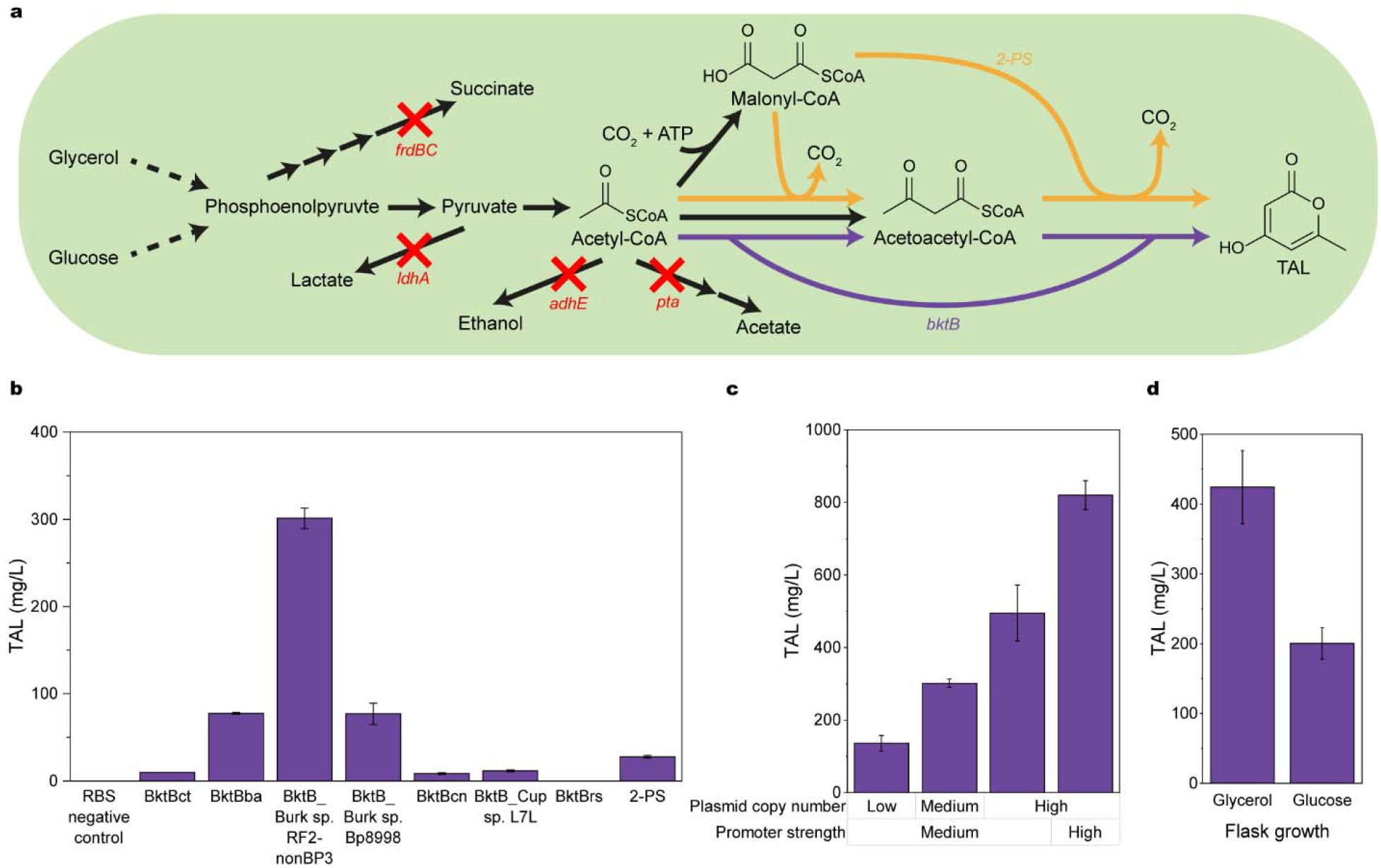
*In vivo* TAL production. a) Pathway scheme of TAL production host strain JBEI-3695, an *E. coli* BW25113 strain with most mixed acid fermentation pathways eliminated through knock-out of *adhE, ldhA, frdBC* and *pta* (indicated by red crosses) for improved acetyl-CoA supply. Black arrows indicate native pathways related to conversion of carbon sources to TAL; purple arrows indicate the reactions catalyzed by overexpressed BktBs; orange arrows mean the reactions catalyzed by overexpressed 2-PS. b) TAL titers of JBEI-3695 expressing different BktBs and 2-PS on pBbA5a, a medium copy number vector with medium strength promoter. c) TAL titers of JBEI-3695 expressing bktBbr on plasmids with different copy numbers and different promoter strengths. Production runs (b and c) were performed in 24-well plates with glycerol as carbon source. d) TAL titers from shake flask growth of JBEI-3695 pBbE1a-bktBbr, which expresses BktBbr on a high copy number vector with a strong promoter showing highest TAL titer in 24-well plates. The experiments were performed in shake flasks with glycerol or glucose carbon source. All growths were conducted under 25 oC.

### Enhanced *E. coli* TAL production in fed-batch fermentation

Fed-batch fermentation for TAL production with *E. coli* using 2-PS has been well studied^17,18,25^. However, fed-batch fermentation with *E. coli* using BktB has not been reported in the literature to our knowledge. Thus, we performed a 1-L fed-batch fermentation of *E. coli* JBEI-3695 pBbA5a-bktBbr with glucose or glycerol as a carbon source. During the 5-day fermentation, cells grew faster with glycerol, while TAL production was higher with glucose. Compared with ∼1.6 g/L TAL production in glycerol media, ∼2.8 g/L TAL was produced in glucose media, becoming the highest titer ever reported using *E. coli* **(Fig. 4a, 4b, Extended Data Fig. 4a, 4b)**. Further analysis of the glucose and secondary metabolite levels during the fermentation indicates that less TAL was produced after 74 h than before 74 h. Although *E. coli* JBEI-3695 had nearly all mixed-acid production enzymes knocked out, the acetic acid was probably produced by the remaining enzyme PoxB ^32^. The acetate was probably used as a carbon source for growth and TAL production after glucose was exhausted **(Extended Data Fig. 4a)**.

**Fig. 4.**
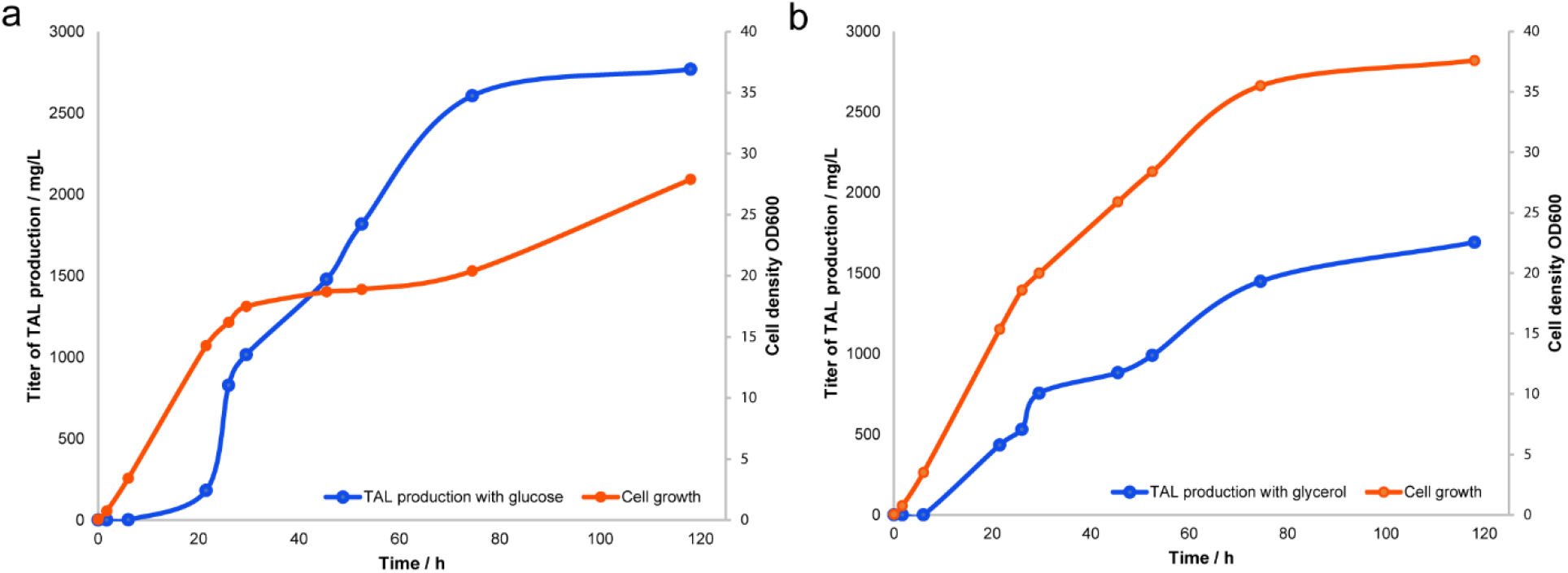
Fed-batch fermentation for enhanced TAL production. *E. coli* JBEI-3695 with pBbA5a-BktBbr was used for TAL production, with a) glucose or b) glycerol as carbon source. The fermentation was performed for 5 days. IPTG was added for protein expression at 4 h. Carbon source feeding was controlled using a DO detector. Samples for measurement of cell density (OD_600_) and TAL titer taken were removed from the bioreactor at different time points and measured.

It is commonly accepted that TAL poses some toxicity to *E. coli*, but there has been no detailed study of this toxicity to our knowledge. Overcoming TAL toxicity is critical for TAL production in *E. coli*, particularly if a very efficient TAL production enzyme, like BktB, is used. Here, three strains were selected to test the TAL toxicity in the TAL production medium with various amounts of TAL spiked into the medium. These three strains were BW25113, JBEI-3695 (BW25113 with deletions of genes *adhE, ldhA, frdBC* and *pta*), and JBEI-3695 pBbE1a-BktBbr which produces TAL (**Extended Data Table 2**). After monitoring cell growth for 78 h at 37 oC, all three strains exhibited comparable TAL toxicity **(Fig. 5a-c)**. Cells harboring a plasmid expressing BktB experienced higher growth inhibition due to potential production of TAL (**Fig. 5c**). Overall, the cell growth inhibition was only mildly inhibited with 2 g/L TAL, while a significant inhibition occurred at 4 g/L TAL, and no cell growth observed with 6 g/L and 8 g/L TAL. This could also explain why the highest titer achieved in bioreactors was not higher than 2.8 g/L. In conclusion, production of TAL in *E. coli* becomes challenging due to such toxicity, although this spike test does not represent the actual fermentation conditions.

**Fig. 5.**
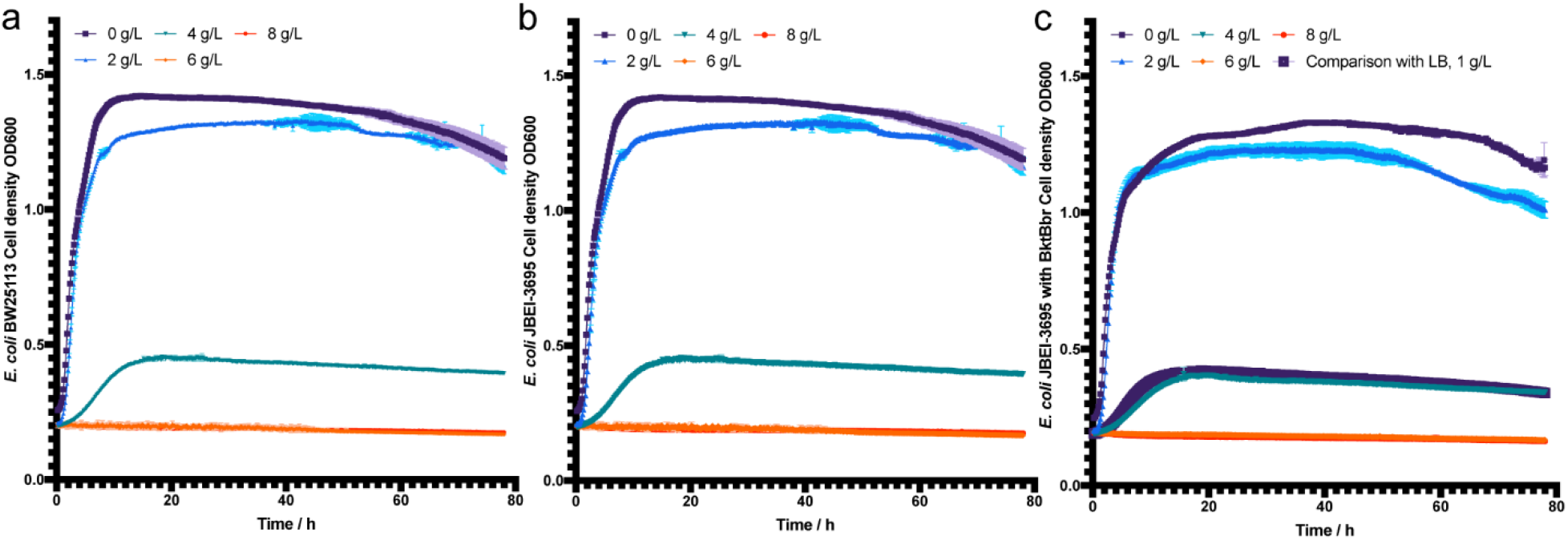
TAL toxicity to various *E. coli* strains. The incubation was conducted in a 48-well plate at 37 [, running for 78 h. Cell growth is monitored at OD_600_. Three biological replicates were included, contributing to the error bars of standard deviation. a) *E. coli* BW25113. b) *E. coli* JBEI-3695. c) *E. coli* JBEI-3695 pBbE1a-BktBbr and hence with TAL production ability.

TAL toxicity is different for *E. coli* in LB broth, which is not as rich as the TAL production medium. At 1 g/L TAL, the toxicity was already so high that the cell growth was significantly inhibited, and no adaptation to TAL toxicity was observed over a 100-h period **(Extended Data Fig. 5)**. In comparison, cell growth inhibition was equivalent with 4 g/L TAL in the TAL production medium and 1 g/L TAL in LB medium (**Fig. 5c**). This finding may lead to a solution to reduce the TAL toxicity for *E. coli* because the richer medium enhances most protein expression levels^33^, possibly including transporters responsible for exporting intracellular TAL from the cell. Identification and up-regulation of the TAL transporters in *E. coli* may be important for it to achieve the high TAL titers produced by yeast. Such transporter engineering, for example efflux pumps, in *E. coli* has been reported to reduce the toxicity of biofuels^34^. Therefore, further increases of TAL production by *E. coli* may require strain engineering or adaptive laboratory evolution to mitigate TAL toxicity^35^. Additionally, an efficient organic overlay could be also helpful by transferring TAL from the fermentation broth to the overlay solution.

## Summary

As a bio-privileged molecule for organic synthesis in industrial manufacturing^15^, TAL has attracted a growing interest. *Yarrowia lipolitica* and *Rhodotorula toruloides* have been engineered to produce 36 g/L and 28 g/L TAL, respectively ^14,25^. However, both studies used 2-PS as the TAL producer. 2-PS consumes intracellular malonyl-CoA, which is critical for cell growth and maintaining other basic cell activities, and whose supply from acetyl-CoA suffers from high energy use to fix carbon dioxide and allosteric feedback. In this scenario, polyketoacyl-CoA thiolase BktB exhibits advantages as an alternative TAL producer by bypassing malonyl-CoA and directly using acetyl-CoA, one of the most abundant intracellular CoA derivatives. In this study, BktBs from *Burkholderia* were shown to be superior for TAL production, compared with the first reported BktB from *C. necator*. Highlighting the featured structures and amino acids found in BktBcn, a phylogenetic analysis enabled us to discover other *Burkholderia* BktBs among the numerous thiolases. These BktBs performed similarly *in vitro* and *in vivo*, and BktBbr led to a record-breaking titer of TAL production, 2.8 g/L, in *E. coli*, even if the production may have been inhibited by TAL toxicity. Undoubtedly, this superior TAL producing enzyme could be used in yeast for TAL production, by enhancing the acetyl-CoA level. Furthermore, due to the large number of secondary metabolites potentially synthesized from TAL^36^, there could be even more efficient BktBs than the best one in this study. These additional enzymes, and their associated structures and functions, would enable us to understand the natural evolution of BktBs and to potentially produce even more TAL using engineered microorganisms. Moreover, more BktBs could be discovered in the future to serve as potential alternatives to native polyketide synthases for biosynthesis of diverse small polyketides.

## Methods

### Strain construction

Strain JBEI-3695 was constructed through step-by-step knockouts of *adhE, ldhA, frdBC* and *pta* in *E. coli* K-12 BW25113. Each step of knock-out was conducted through λ-Red-mediated recombination based replacement of target gene with kanamycin resistance marker amplified from corresponding strains of Keio Collection, a library of *E. coli* strains with each single gene replaced with kanamycin resistance^37^, followed by FLP-mediated removal of resistance^38^. All strains were stored in 30% glycerol under -80 oC. Strains were streaked or plated on LB agarose plates (Teknova, Hollister, CA) with addition of antibiotics at the following concentrations if necessary: carbenicllin (100 μg/ mL) and kanamycin (50 μg/ mL). Unless specified, all media components and supplemented chemicals for strain growths and constructions were acquired from Sigma-Aldrich (St. Louis, MO), Thermo Fisher Scientific (Waltham, MA) and Becton, Dickinson and Company (Franklin Lakes, NJ).

### Plasmid construction

All genes encoding BktBs and 2-PS were synthesized with GenSmart™ codon optimization, based on the “Population Immune Algorithm”, for *E. coli* and subcloned into pET-28a(+)-TEV vector between NdeI/XhoI restriction sites by Genscript (Piscataway, NJ). Genbank Accession Numbers of proteins tested in this study can be seen in **Extended Data Table 1**, and the list of plasmids used in this study can be seen in **Extended Data Table 2**. Other plasmids were constructed through Gibson Assembly between vector and genetic parts amplified by PCR with Phusion High-Fidelity DNA polymerase with all kits from New England Biolabs (Ipswich, MA). The PCR amplified DNAs were purified by DNA Clean and Concentrator or Zymoclean™ Gel DNA Recovery Kit (Zymo Research, Irvine, CA). Primers were synthesized by Integrated DNA Technologies (Coralville, IA). Biobrick plasmids^31^ with different copy numbers and promoter strengths were used as vectors. The constructed plasmids were chemically transformed to DH5a competent cells (New England Biolabs) and then their sequences were confirmed through Sanger Sequencing by Sequetech Corporation (Mountain View, CA). The sequence confirmed plasmids were then transformed into JBEI-3695 or other host strains through TSS chemical transformation.

### Protein purification

*E. coli* BL21(DE3) transformed with pET-28a (+)-TEV harboring the gene encoding BktBs or 2-PS was inoculated into 10 mL of LB broth containing kanamycin, incubated at 37 □ overnight at 225 rpm shaking. This seed culture was used to inoculate 1 L of LB broth containing kanamycin and then incubated at 37 □ for 2 to 3 hours at 225 rpm shaking. When the OD_600_ reached 0.6, the culture was cooled on ice for 15∼20 min. 0.2 mM isopropyl β-D-1-thiogalactopyranoside (IPTG) was added to induce protein expression followed by incubation at 18 □ to allow the protein expression. Cells were harvested by centrifugation at 5,000 × g at 4 □. Harvested cells were transferred to a 50 mL Falcon tube kept at -80□ until needed. Pellets from 0.2 L of cell culture were thawed at room temperature and resuspended in a 10 mL cell lysis buffer (50 mM sodium phosphate, pH 7.4, 300 mM NaCl, 10 % glycerol). Cells were lysed by sonication for 15 min (500-Watt, 30% amplitude, pulse 5s on and 10s off). Cell debris and other insoluble material were removed by centrifugation for 45 min at 13,000 × g at 4 □. The soluble fraction was filtered using a 0.22 μm filter, mixed with 1 mL of Ni-NTA resin (Thermo Scientific, USA), and incubated at 4 □ for 1 hour under constant mixing. The resulting resin was washed twice with 10 mL wash buffer (50mM sodium phosphate, pH 7.4, 0.3 M NaCl, 20 mM imidazole), and the target protein was eluted using 5 mL elution buffer (50 mM sodium phosphate, pH 7.4, 0.3 M NaCl, 200mM imidazole). Purity of the target proteins was checked using an SDS-PAGE gel. The protein concentration was quantified by Bradford Assays using the BSA standard curve. Purified proteins were kept at -80 □ in small aliquots until needed.

### Protein sequence alignment and phylogenetic tree construction

Alignment of the protein sequences obtained from Uniprot (https://www.uniprot.org/) were performed in MAFTT (https://mafft.cbrc.jp/alignment/server/) using UniRef50. The aligned sequences were analyzed on ClustalW2 Phylogeny of EMBL-EBI (www.ebi.ac.uk/Tools/phylogeny/) by applying neighbor-joining clustering methods. Visualization of the phylogenetic tree was generated by iTOL (itol.embl.de/tree/). The mode of the tree is circular with rotation of 210 ° and arc of 350 °. The branch lengths were used to represent the circular tree.

### *In vitro* assays

The *in vitro* assays for functional determination of TAL formation by LC-MS were performed in a 200 μl total reaction volume containing 3 mM EDTA, 1 mM acetoacetyl-CoA, 1 mM acetyl-CoA and 50 μg (∼6 μM) of purified enzymes in 100 mM potassium phosphate buffer, pH 7.2. The reaction is incubated at room temperature for 30 min before LC-MS analysis for TAL synthesis.

The *in vitro* assays for kinetics using α-KGDH for CoA conversion has been reported ^39^. For BktBs, enzyme activities were performed at 22 □ in a 100 μL total reaction volume containing 50 mM sodium phosphate, 3 mM EDTA, 1mM dithiothreitol, 24 mU/μL α-ketoglutarate dehydrogenase, 2.5 mM NAD^+^, 0.4 mM thiamine pyrophosphate, 2 mM α-ketoglutaric acid, 100 μM acetoacetyl-CoA, 100 μM acetyl-CoA, and 3 μM of purified enzyme. The increase in absorbance at 460 nm (NADH generation) was monitored in a SpectraMax M2 (Molecular Devices, USA) microplate reader for 30 min. Data points in the initial 4 min were used to generate the linear curve for the quantification of the initial reaction rate *v*, slope of the linear curve.

### *In vivo* TAL production

The cell growth medium for TAL production was EZ-rich media (Teknova, Hollister, CA) supplemented with 10 g/L tryptone, 5 g/L yeast extract, 5 mM calcium pantothenate and 20 g/L glycerol or glucose carbon source and with replacement of 1.32 mM K_2_HPO_4_ to 2.8 mM Na_2_HPO_4_ when glycerol was the carbon source, modified from a previous report^40^. During comparisons of different BktBs, cells were grown in 2.5 mL media contained in Axygen® 24-well Clear V-Bottom 10 mL Polypropylene Rectangular Well Deep Well Plates (Corning, Corning, NY) sealed with sterile AeraSeal™ films (Excel Scientific, Victorville, CA), while the flask growth was performed in 20 mL media contained in 250 mL Pyrex narrow mouth Erlenmeyer flasks with screw cap (Corning). Before cell growth, a small portion of glycerol stock was inoculated into a glass tube containing 5 mL LB with carbenicllin and grown overnight under 37 oC. 1% volume of overnight seed cultures were then inoculated in the media followed by incubation under 37 oC initially. After three to four hours when cell growth reached exponential phase, 100 μM IPTG was added for induction and cells were transferred to Kuhner ISF-1-W incubators (Kuhner Shaker, Birsfelden, Switzerland) set at 200 rpm shaking and different temperatures of 37, 30, 28 and 25 oC as needed. Cell cultures were collected for analysis after 48 hours of post-induction growth. At least three biological replicates were grown in each round, and pre-screening of the colony with highest titer was performed if colony-to-colony variance was high.

### LC-MS TAL identification and quantification

TAL analysis was performed using an Agilent LC/MSD IQ system (Agilent Technologies, Santa Clara, USA). Both *in vitro* and *in vivo*, 50∼100 μL samples were mixed with equal volume of 100 % acetonitrile containing internal standard, (5S,6S)-6-isopropyl-5-methyldihydro-2H-pyran-2,4(3H)-dione (C_9_H_14_O_3_, 170.21 Da), followed by filtration with 3 MWCO 96-well plate (Pall Corporation, Port Washington, NY). Kinetex XB-C18, 2.6 μm, 3 mm x 100 mm column (Phenomenex, USA) was used for LC separation of the molecules. 3 μL sample was injected in the LC-MS for analysis. Mobile phase A was water with 0.1 % formic acid, and mobile phase B was methanol with 0.1 % formic acid. The detailed method and parameters are shown in **Extended Data Table 3**. MS was run in negative mode, leading to m/z of TAL is 125.1 and m/z of internal standard is 169.1. Target peaks in SIM mode were auto integrated for peak area. The peak area of TAL is calibrated by the internal standard signal.

### HPLC TAL quantification

The TAL titers were quantified using an Agilent 1200 HPLC System (Agilent Technologies, Santa Clara, CA) with Diode Array Detector (DAD) under 298 nm. 500 μL of collected TAL production broth was mixed with 500 μL acetonitrile, followed by 5000 × g centrifuge for 5 min. 300 μL of supernatant was filtered under 3 kDa through Amicon Ultra-0.5 mL Centrifugal Filters (Millipore, Burlington, MA) or 96-well plates (Pall Corporation, Port Washington, NY), then transferred to HPLC vials. Different concentrations of TAL standard samples were also prepared and measured through the same way to create the standard curve. Kinetex 2.6 μm EVO C18 100 Å LC Column with 100 × 4.6 mm size (Phenomenex, Torrance, CA) was used. The sample injection volume was 10 μL. The mobile phase and LC program were the same as LC-MS.

### Fed-batch fermentation conditions

Fed-batch bioreactor experiments were performed in 2-L bench top glass fermenters (Biostat B, Sartorius Stedim, Göttingen, Germany) equipped with two 6-blade Rushton impellers. Cells were grown at 37 °C, 200 rpm for 16 h in LB media. All tanks were batched with 1 L of LB-like EZ-Rich media containing glucose or glycerol as carbon source. The bioreactors were inoculated with seed cultures at a starting OD of 0.05. Temperature, agitation, and air flow were maintained constant at 22 °C, 300 rpm and 0.5 vvm, respectively and pH was controlled to 7.0 using 10% (v/v) H_2_SO_4_ and 14% (v/v) NH_4_OH. Protein expression was induced with 0.1 mM IPTG when OD reached 0.6 and temperature was adjusted to 22 °C after induction. Fed-batch experiments employed a DO signal-triggered feeding loop (ΔDO = 15 %, Flow rate = 40mL/h, Pump duration = 5 min). Glucose feeding started when the initial amount of glucose was depleted with a feed solution containing 600 g/L glucose or glycerol and 50 mg/L carbenicillin. 2 ml samples were taken in regular intervals and centrifuged at 15,000 × g for 5 min. For analysis, the supernatant was filtered (0.2 mm) and stored at -20 °C, and the cell pellet was stored at -80 °C.

## Extended Data Figures and Tables

**Extended Data Table 1.**
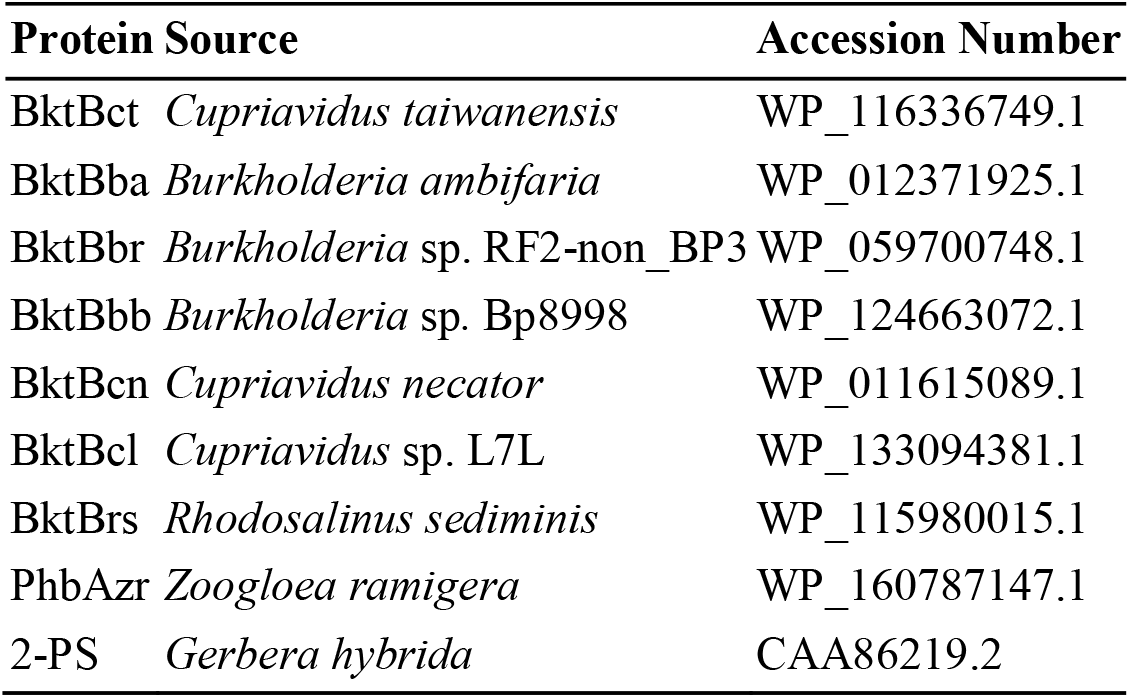
GenBank Accession numbers of proteins used in this study.

**Extended Data Table 2.**
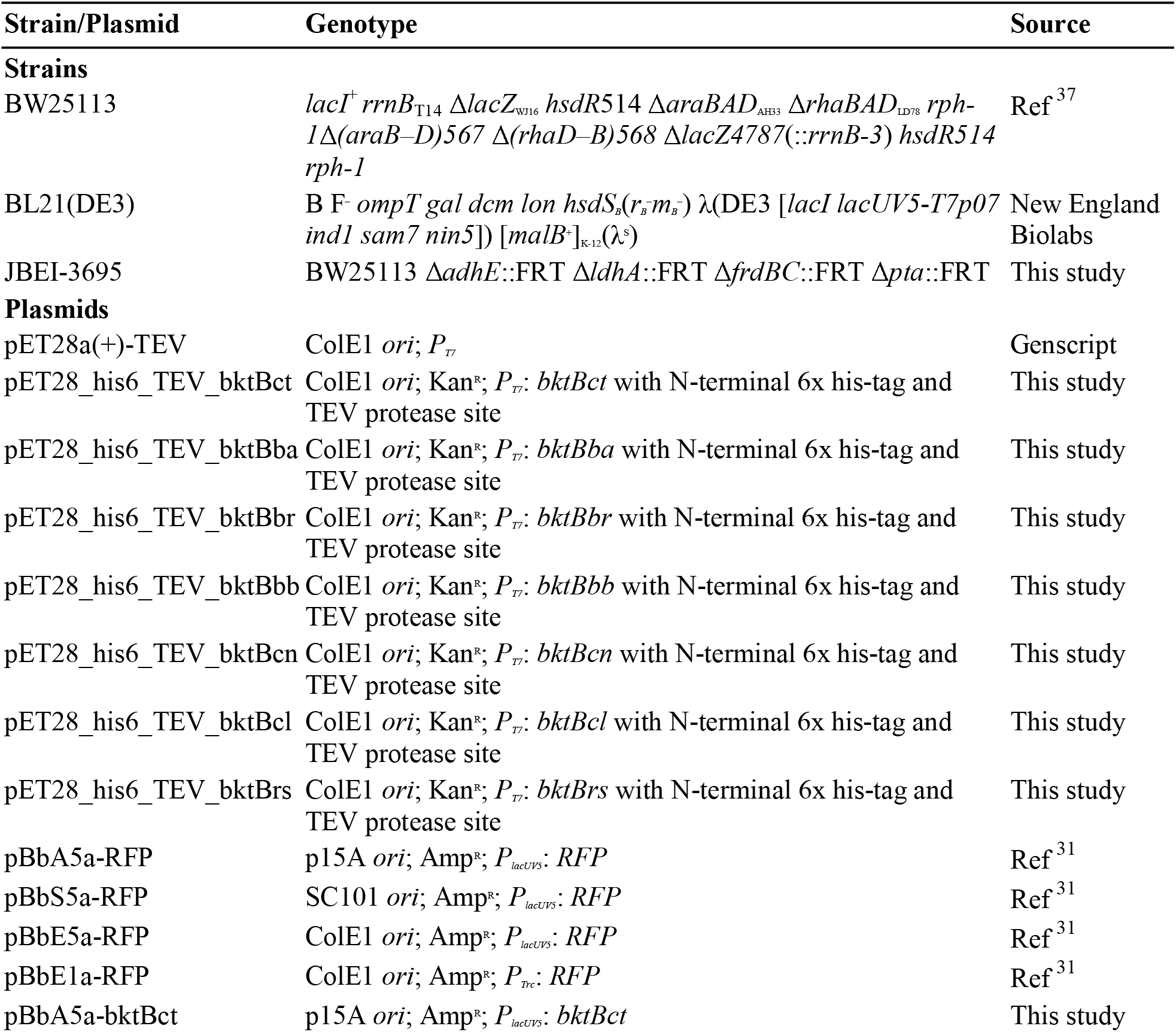
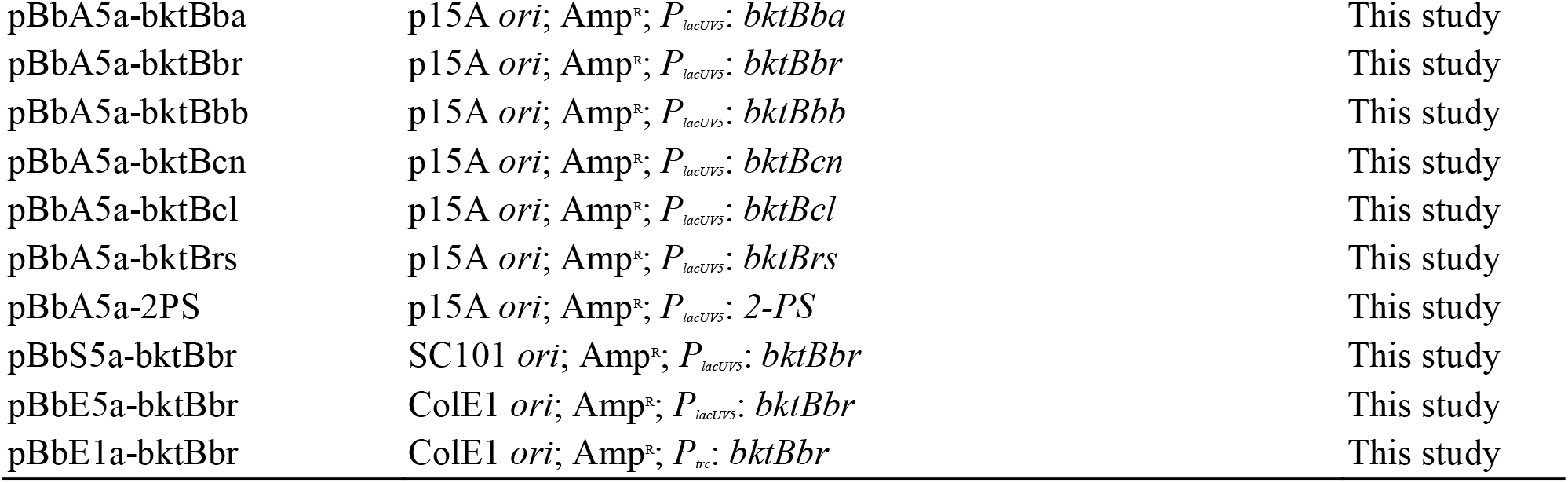
Strains and plasmids used in this study.

**Extended Data Table 3.**
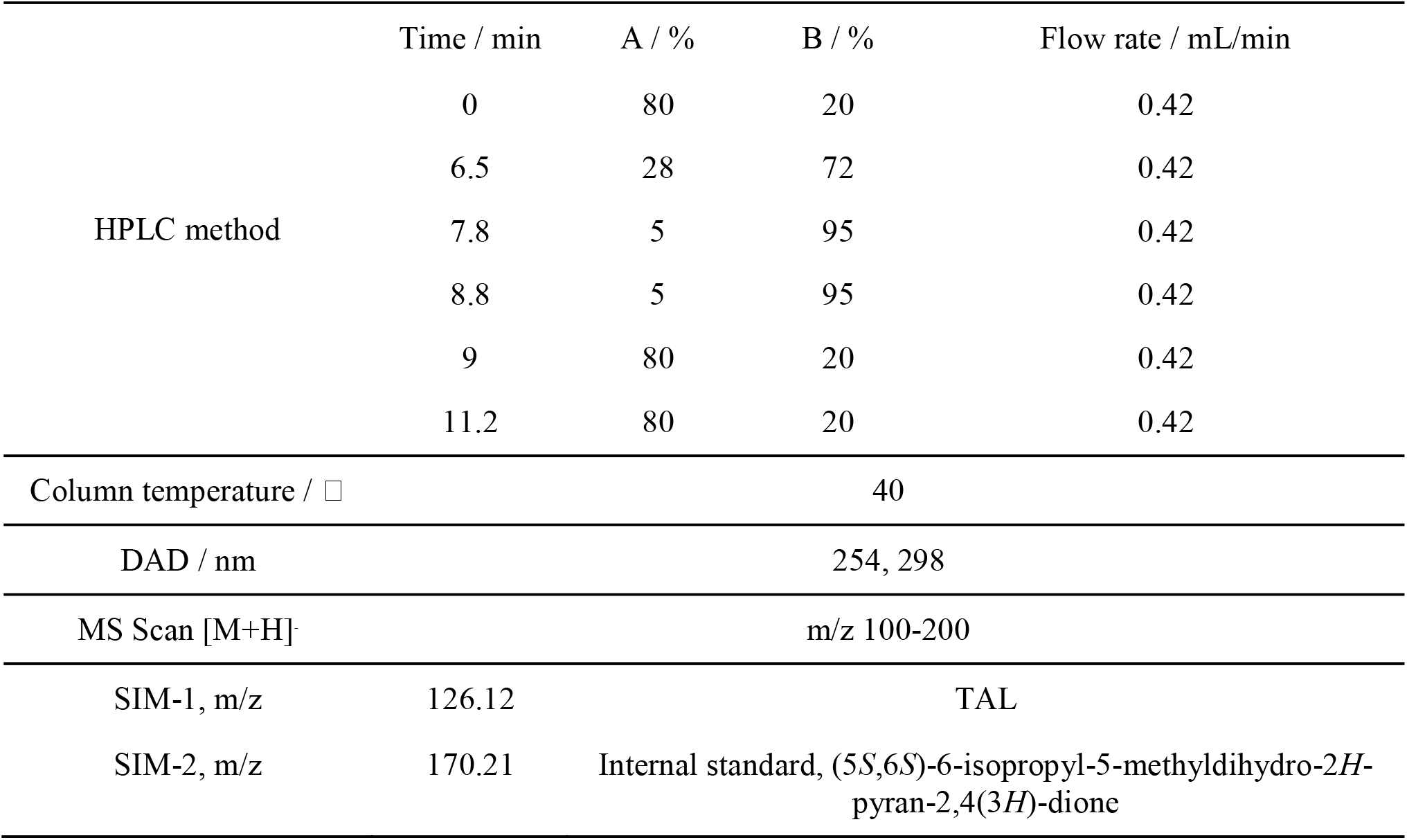
LC-MS method and parameters for TAL identification and quantification.

|  | Time / min | A / % | B / % | Flow rate / mL/min |
| --- | --- | --- | --- | --- |
| HPLC method | 0 | 80 | 20 | 0.42 |
|  | 6.5 | 28 | 72 | 0.42 |
|  | 7.8 | 5 | 95 | 0.42 |
|  | 8.8 | 5 | 95 | 0.42 |
|  | 9 | 80 | 20 | 0.42 |
|  | 11.2 | 80 | 20 | 0.42 |
| Column temperature / °C |  | 40 |  |  |
| DAD / nm |  | 254, 298 |  |  |
| MS Scan [M+H]<sup>+</sup> |  | m/z 100-200 |  |  |
| SIM-1, m/z | 126.12 | TAL |  |  |
| SIM-2, m/z | 170.21 | Internal standard, (5*S*,6*S*)-6-isopropyl-5-methyldihydro-2*H*-pyran-2,4(3*H*)-dione |  |  |

**Extended Data Figure 1.**
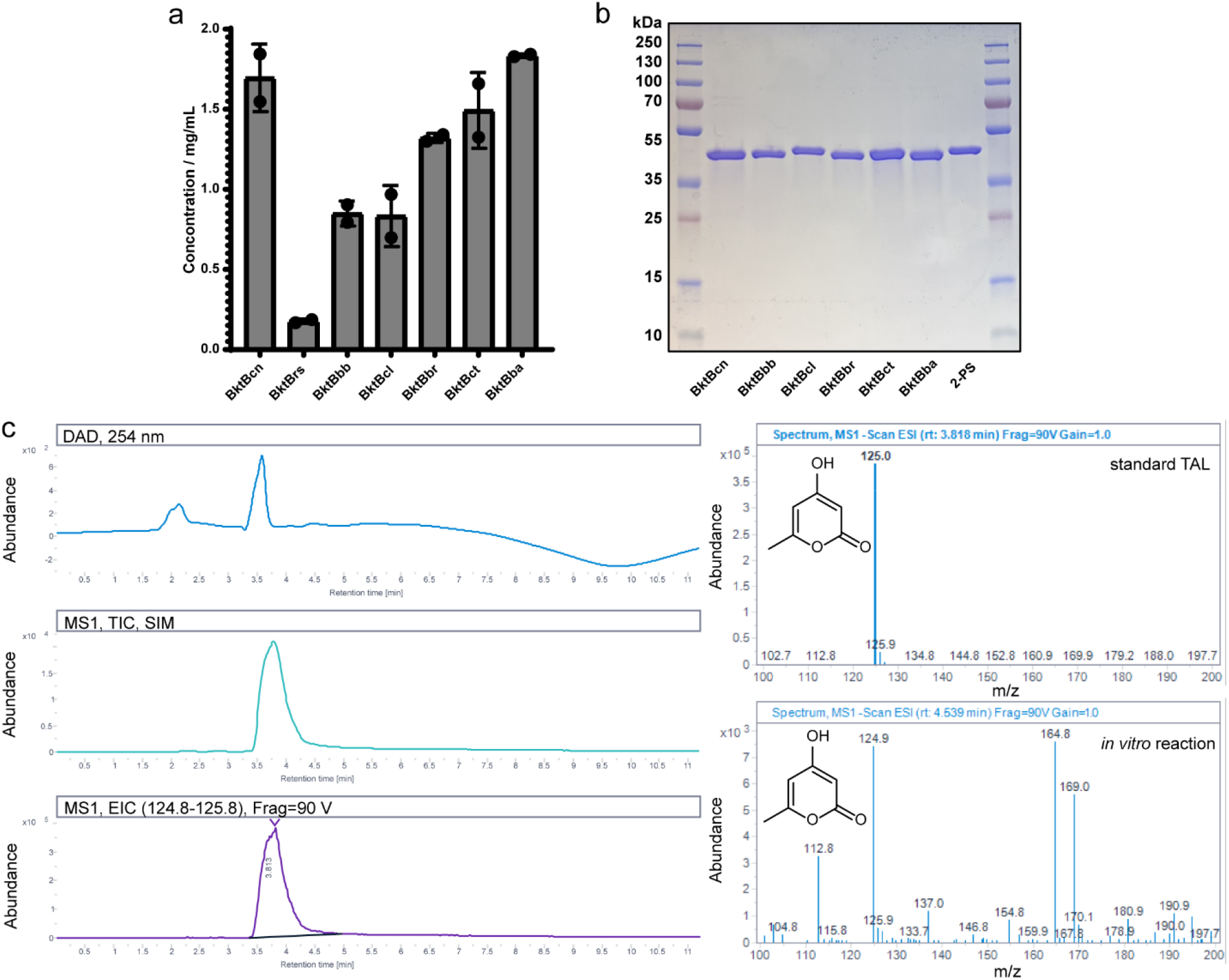
Purified proteins and LC-MS of *in vitro* TAL formation. a) Protein concentrations, measured by Bradford assay, of purified BktBs from *E. coli* BL21(DE3) by nickel resin. b) SDS-PAGE gel result of purified proteins used in this study. 55 μg of proteins was loaded into each lane. SDS-PAGE gel (Bio-Rad, USA) was precast to 4∼20 %, run at 140 V for 1 h. c) LC-MS result of TAL identification with mass spectrum. TAL eluted at 3.82 min. Mass spectrum of pure TAL standard and *in vitro* reaction of TAL production are shown on the right. m/z of 124.9 is TAL, and 169.0 is the internal standard.

**Extended Data Figure 2.**
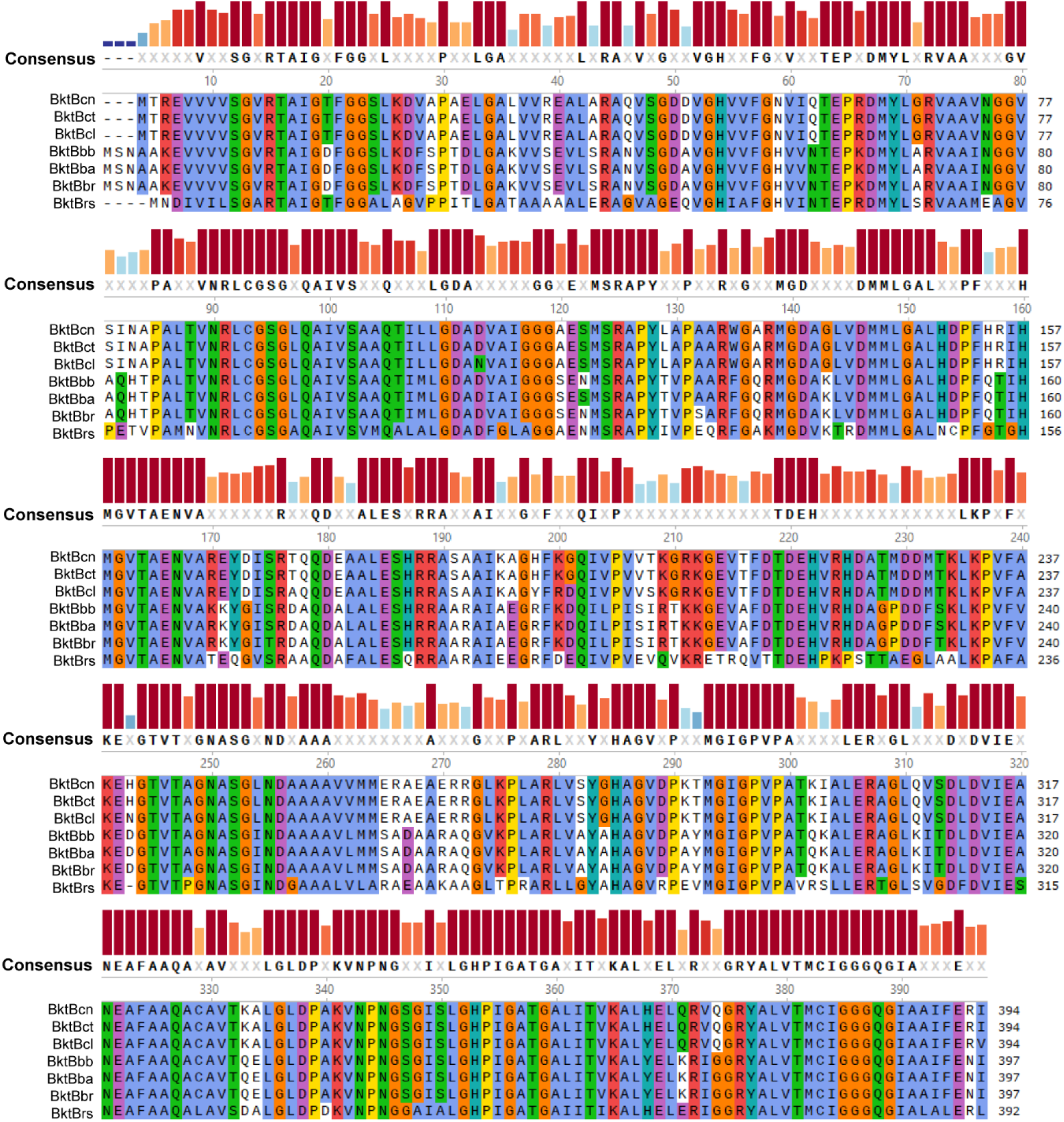
Complete protein sequence alignment of BktBs. Seven selected BktBs are aligned by MAFFT. The consensus amino acid residues are highlighted in different colors and shown in the Consensus row. The bar graph shown above each sequence alignment represents the identity % of amino acids at different positions.

**Extended Data Figure 3.**
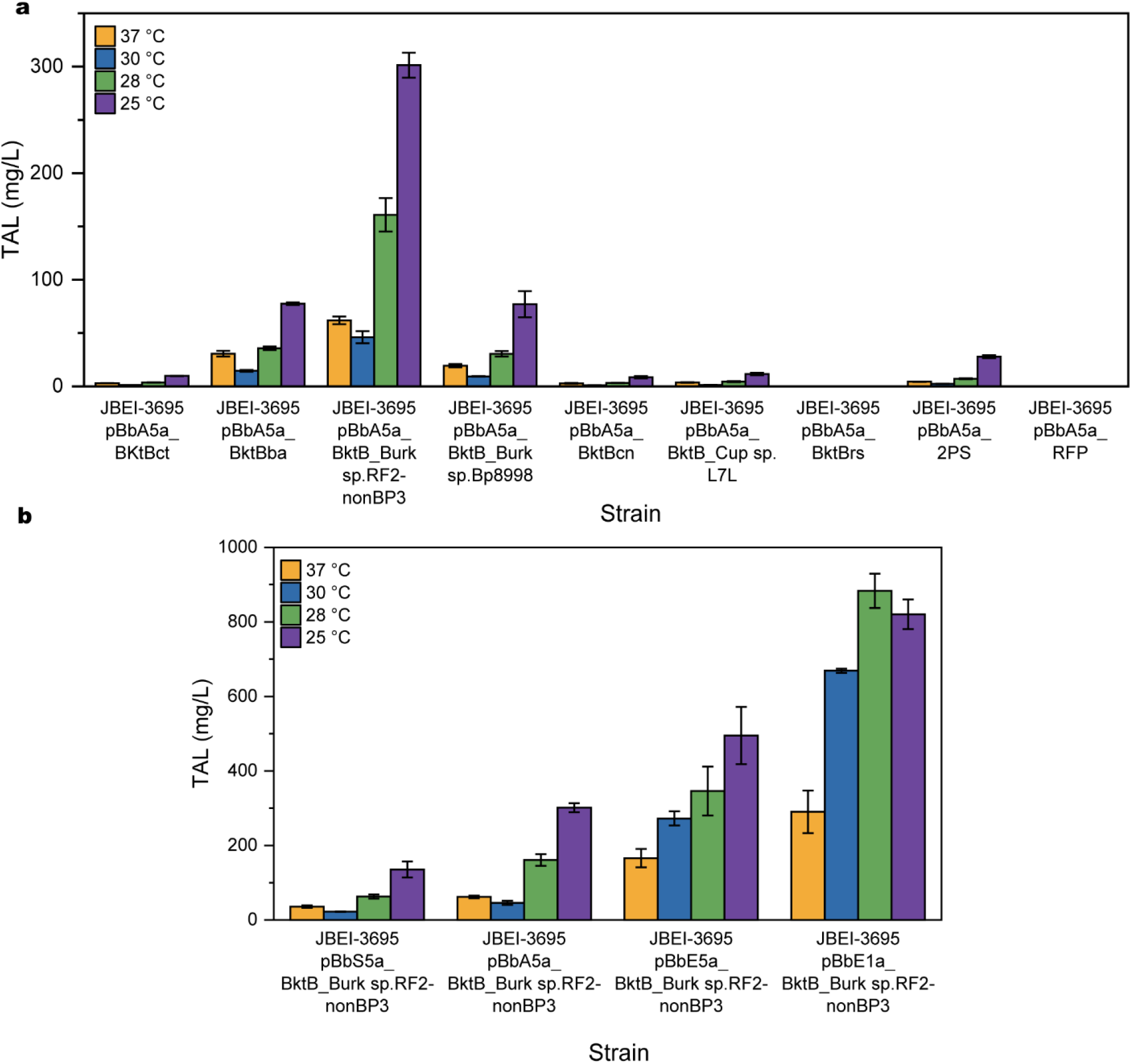
*In vivo* TAL production by *E. coli* strains expressing different BktBs and 2-PS in 24-well plates under different temperatures. The host strain was JBEI-3695 (BW25113 Δ*adhE* Δ*ldhA* Δ*frdBC* Δ*pta*), and strains were cultivated with glycerol carbon source.

**Extended Data Figure 4.**
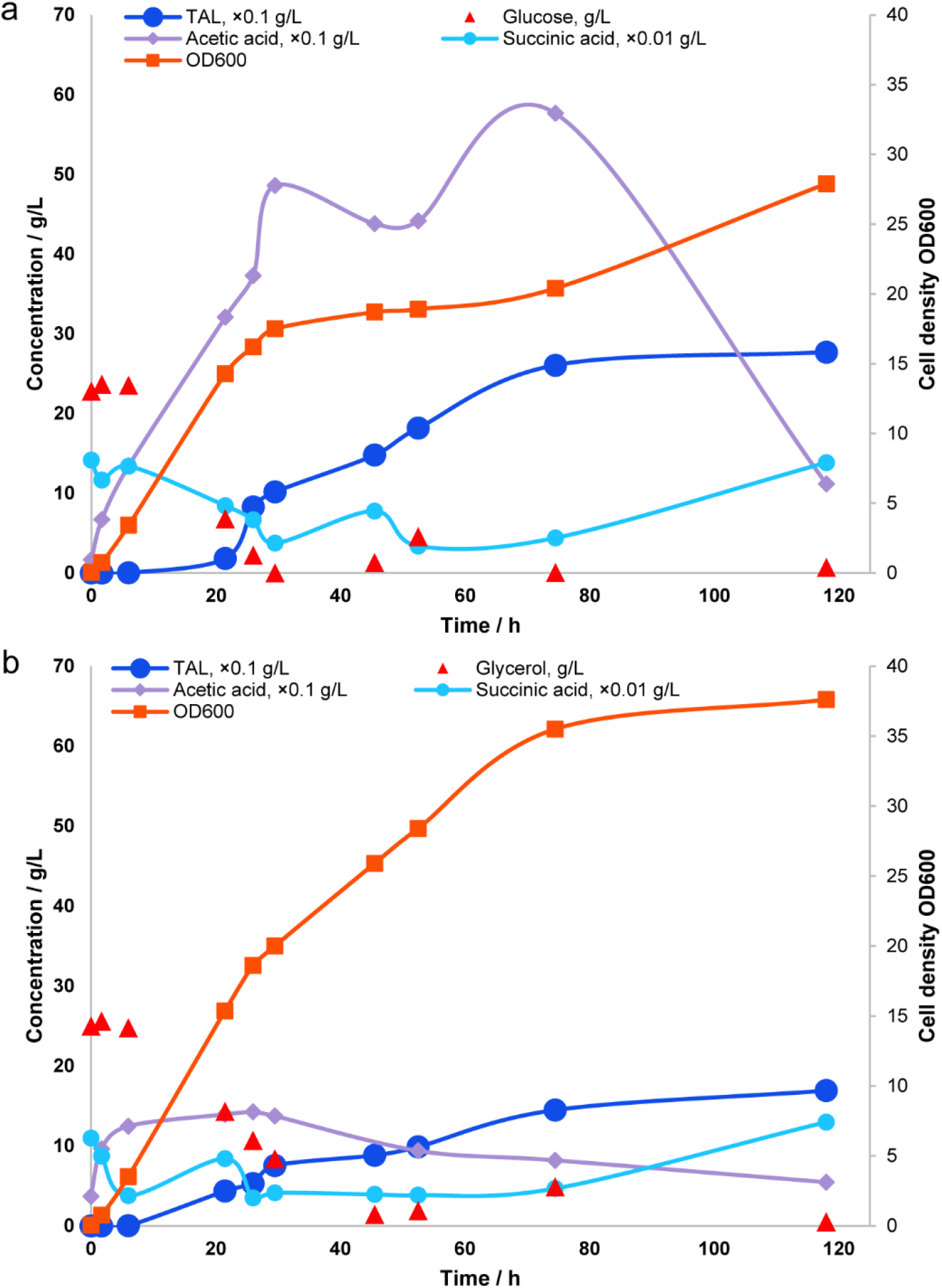
Analysis of 1 L fed-batch fermentation with *E. coli* JBEI-3695. Samples were collected at different time points, allowing the HPLC analysis of the glucose / glycerol level and metabolites levels. a) Growth and production with glucose in the medium. b) Growth and production with glycerol in the medium. The y axis is adjusted to be the same between a) and b).

**Extended Data Figure 5.**
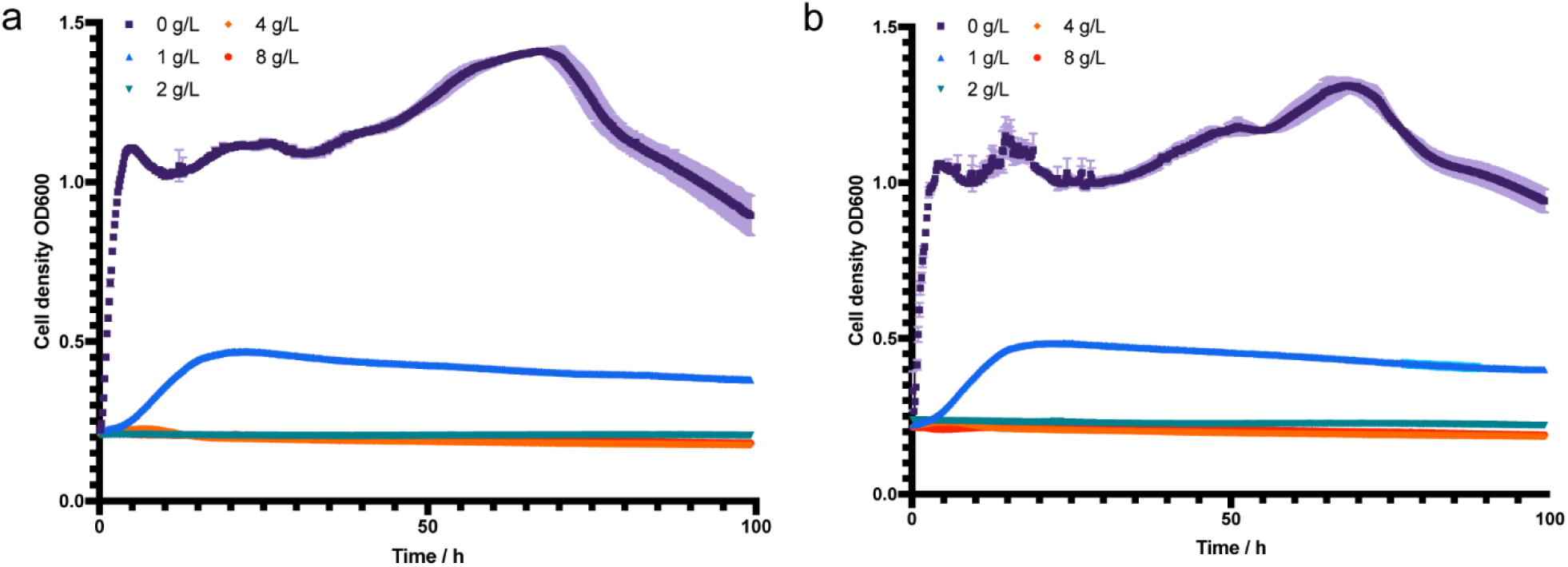
TAL toxicity to *E. coli* strains in LB medium. The incubation was performed in a 48-well plate under 37 □ for 100 h. Cell growth was monitored by measuring OD600. Three biological replicates were included, contributing to the error bars of standard deviation. a) *E. coli* BW25113, b) *E. coli* JBEI-3695.

